# Physical fitness and motor competence performance characteristics of elite Chinese youth athletes from four track and field throwing disciplines – a cross sectional study

**DOI:** 10.1101/2023.07.20.549861

**Authors:** Kewei Zhao, Maximilian Siener, Yifan Zhao, Andreas Hohmann

**Affiliations:** High Performance Research Center, China Institute of Sport Science, Beijing, China; BaySpo–Bayreuth Center of Sport Science, University of Bayreuth, Bayreuth, Germany; Institute of Professional Sport Education and Sport Qualifications, German Sport University, Cologne, Germany

**Keywords:** shot put, hammer throw, discus throw, javelin throw, discriminant analysis, multilayer perceptron

## Abstract

**Purpose:** Systematic athletic training during adolescence may facilitate the development of sport-specific skills and the expression of sport-specific physical characteristics in young athletes. The aim of this study was to differentiate male athletes aged 14-17 years from four different throwing disciplines using anthropomorphic measurements and motor competence performance tests, in order to test whether athletes from different disciplines have physical form and fitness characteristics consistent with the sport-specific demands of each throwing discipline.

**Methods:** The sample consisted of 289 male youth athletes belonging to the four different throwing disciplines: shot put (n=101), hammer throw (n=16), discus throw (n=63), and javelin throw (n=109). The performance diagnosis comprised three anthropometric measurements, and twelve physical fitness tests. Discriminant analysis and neural network (Multilayer Perceptron) were used to test whether it is possible to discriminate between athletes of the four sports.

**Results:** The results of this study show that for male throwing athletes aged 14-17 years, differences in generic anthropometric and sport performance tests distinguish the talent of more than two-thirds of young athletes based on individual sport (DA: 68.7%; MLP: 72.2%), regardless of the classification method used.

**Conclusion:** The relevance of the three anthropometric parameters and twelve physical fitness measures for talent identification and training monitoring in the track and field throwing disciplines was confirmed. The discus throwers had a height advantage, the shot putters and hammer throwers had better arm strength, while the javelin throwers had better explosive strength and sprint speed. All events, except the hammer throwers, showed a high level of explosive power in the medicine ball forward or backward throw test. This was particularly important for the shot put and discus athletes.

## Introduction

Participation in elite sport training at a young age is associated with the selection of athletes with specific performance prerequisites of a particular sport(1), as well as with the proper development of such features through a systematic training process. For example, athletes in shot put, hammer and discus throw exhibit specific anthropometric characteristics, such as a higher body weight and lean body mass compared to javelin throw athletes(2). Additionally, Carter, et al.(3) and Morrow, et al.(4)found that shot put and discus throw athletes tend to have a superior body height compared to javelin throwers.

In regard to physical fitness, maximum arm and leg strength and ballistic power of the extremities are important performance characteristics for athletes in all four throwing disciplines, as the strength of the arm and leg extension muscles plays an important role in transferring a high momentum to the throwing device(5). Accordingly, Terzis, et al.(6) and Bouhlel, et al.(7) found out that, in shot-put and javelin throw respectively, both maximal leg and arm strength were significantly correlated with throwing performance. Furthermore, this important relationship has been demonstrated not only in comparably heterogeneous athletes at the collegiate level(8,9), but also at the more homogeneous elite performance level(10). In addition to explosiveness, sprinting speed also has a significant impact on the throwing performance, e.g in collegiate shot put athletes, as was found by Caughey and Thomas(11). Finally, core stability(12,13) and flexibility(14) of the athletes play an important role in the throwing disciplines which is not only relevant in the shoulder, but also in the trunk, hip and leg. Although there are no reports on the validity of running endurance tests in track and field throwing disciplines, these tests using basic exercise modes are common in the seasonal preparation phases of all four throwing disciplines. For this reason, it also seems reasonable to consider and test endurance capabilities.

On the basis of such findings, the most relevant sports-specific performance characteristics of the four track and field throw disciplines shot put, hammer throw, discus throw, and javelin throw can be used for talent identification or talent transfer purposes(15,16), as well as for the planning of the athletes’ long-term education(17), and the monitoring of the seasonal training process of youth athletes(18–20). The need to investigate not only differences, but also similarities in the performance characteristics in the four track and field throw disciplines is underlined by the findings of Horst, et al.(21), who could identify identical patterns of individual performance characteristics in shot put, discus, and javelin throwing decathlon athletes.

In long-term talent development programs, talent identification procedures include morphological measures, as well as motor tests data. Accordingly, several talent identification programs in elite sport schools have implemented morphological and physical fitness diagnostics(22–25) to select or transfer young athletes into specific sports, such as alpine skiing(26), soccer(27), tennis (28,29), table tennis (30,31). Talents in particular sport disciplines exhibit a specific make-up of natural abilities (nature) and well-developed performance prerequisites (nurture)(32). Therefore, the predictive validity of such talent characteristics is of paramount importance in the identification and development of promising youth athletes. Although some academics warn against talent identification procedures that are conducted too early(33), these procedures targeted at juveniles are worthwhile for the purpose of helping sports federations to focus their resources on the most talented young athletes(34,35).

Many sports are based on a complex, multi-dimensional performance profile(36). Thus, the talent selection should focus on a multifaceted variety of general physical, physiological, psychomotor, and psychological performance diagnostics(37,38). In general, there is a lack of research investigating the discriminative value of different performance prerequisites across a range of different sport disciplines. However, there have been promising attempts to discriminate between different sports based on their sport-specific performance requirement profiles. For example, Leone et al.(39) were able to discriminate 88% of athletes from four different sports (figure skating, swimming, tennis, and volleyball) using a discriminant analysis that included anthropometric and motor characteristics. Opstoel et al.(40) reported a correct classification of 85.2% of highly active U12 athletes into their own sport (ball sports, dance, gymnastics, martial arts, racket sports, and swimming). In addition, Pion et al. (32) were able to classify 96.4% of 141 Flemish adolescent athletes into nine different sports. In line with our research focus, the findings of Pion et al.(41) in elite male U18 athletes were very promising, as the researchers found a 100% correct classification within the more interrelated martial arts disciplines of judo, karate, and taekwondo. In contrast to the aforementioned studies, discriminant analysis is less accurate when a hold-out of one case (*n*=1) is used to classify on the basis of the discriminant functions obtained from all other cases (n-1). Using this cross-validation strategy in a discriminant analysis with 56 youth athletes aged 12-16 years from six different sports (basketball, fencing, judo, swimming, table tennis, volleyball), Zhao, et al.(25) reported a correct classification of 71.3% of the athletes. Using alternatively the neural network method multilayer perceptron (MLP), and applying a 10% holdout strategy, the authors reported an almost identical classification rate of 71.0%.

Thus, reliable and valid information regarding the potential of talented athletes in specific sports based on morphological parameters, motor abilities and skills, and physiological diagnostics is a valuable tool in performance development programs for clubs and sport federations. The main reasons for scientific uncertainties in talent orientation arise from the often-undifferentiated mixture of general as well as sport-specific tests in talent identification campaigns. The unsystematic timing of cross-sectional diagnostics at single points in time during the long-term athletic development process also contributes to the aforementioned uncertainty. Thus, it is not surprising that the great variety of study design parameters have led to inconsistent research results, providing an inconsistent picture with regard to the discriminative validity of talent features addressing general and sport-specific performance prerequisites. Therefore, for talent identification and development, there is a need for multifaceted test batteries that can distinguish between the anthropometric and specific fitness attributes necessary for different sports.

Although the prediction of long-term success is still debatable, the talent orientation method of recommending suitable sports to young athletes according to their individual talent make-up seems feasible in early childhood(32), as well as part of a talent transfer strategy at older ages(42). Thus, the general purpose of this study is to discriminate elite male athletes from four different track and field throwing disciplines using morphological and motor performance tests. Additionally, it will be investigated whether the athletes from the different disciplines have a sport-specific anthropometric and motor performance profile that corresponds to the specific requirements of the respective throwing discipline. It was hypothesized that a generic test battery would have sufficient discriminative validity to assign athletes to their own sport based on their individual profile of test scores. In this case, the athletic performance prerequisites could serve as a scientific knowledge background for talent identification, athlete training, or talent transfer in this field of athletics.

## Materials and Methods

### General Study Design

In this study, Chinese male junior athletes from four different track and field throwing disciplines were tested for their physical fitness and motor competence using a generic test battery. In addition, body height and body weight were collected from each participant. Subsequently, it was examined whether the test results were directly related to the achieved throwing performances and, with the help of linear and non-linear classification analyses, it was analyzed whether a discrimination of the sport disciplines is possible on the basis of the test values.

### Participants

A sample of *N* = 289 male youth elite track and field athletes aged 14 to 18 years (*M* = 15.94 yrs; *SD* = 1.02) from more than 30 municipal elite sports schools from all-over China were selected for this study. The track and field athletes practiced in one of the four different throwing disciplines shot put (n=101), hammer throw (n=16), discus throw (n=63), and javelin throw (n=109), and all participated in provincial and/or national competitions for junior athletes in China. All athletes trained at least twice a day for a total of 18 hours over six training days a week. All athletes were performing at a high level in their respective sport, representing their provinces in nation-wide competitions. The participants were recruited according to the ethical standards of the China Institute of Sport Science (CISS), and the test battery and the test execution were organized by the CISS.

This study was conducted in accordance with the recommendations of the “Science Research Ethics Committee at the China Institute of Sport Science (CISS)” with written informed consent from all subjects. Ethics approval and parental written informed consent was obtained from the participants in this study in accordance with the declaration of Helsinki. Parents of all athletes were informed of the study protocol, which was outlined in an information letter. No data collection took place without parents’ consent.

#### Measurements

The participants completed two morphological, and twelve motor tests, which were administered by expert sports school staff. All tests were completed on the same day, with the Chief Judge video-linked to the referees at each test location to start each event in unison according to the content of the test. The tests started at 9 a.m., and all athletes refrained from strenuous exercise the day before the test session.

### Throw Performances

The discipline-specific throwing performances of the four groups of athletes were measured using four discipline-specific standing throws (standing shot put, spin hammer throw without body rotation, standing discus, standing javelin throw). All four measurements were performed in the original competition setting in the municipal stadiums of the respective school city, and were carried out by officially licensed referees.

### Morphological Characteristics

Body height (BH) to the nearest 0.1 cm (Height Tester, Donghuateng Sports Apparatus Ltd, Beijing, China), and body weight (BW) to the nearest 0.1 kg (calibrated Seca Alpha 770) were measured according to standardized test procedures (43,44), and Body Mass Index (BMI) was calculated. As in some cases only the BMI was reported instead of the BH and BW data, in most of the analyses the BMI was used.

### Motor Characteristics

The physical fitness (PF) of the participants was assessed by nine generic test procedures. The PF test battery diagnosed maximal explosive leg strength (measured by two trials of a standing long jump and triple step jump without run-up), arm strength (measured by a single trial of pull-ups with chin to the bar, and two trials of a 2-kg medicine ball throw forward and backward (starting position: both feet on the ground behind a line; no step or run-up; measurement perpendicular to measuring tape), core strength (measured by a single trial of the plank test), endurance (measured by a 2-km run on the athletics track), and running speed (measured by single trials of the 30-m and 60-m running sprint with light-gates, Brower Timing Systems; Draper, USA; starting position was 0.3 m behind the starting line). The motor competence (MC) of the participants was diagnosed by three generic tests for flexibility (measured by two trials of the sit and reach test), agility (measured by the total result of three trials of Hexagonal jumping), and balance (measured by a single trial of a one-legged stance on a balance pad (balance pad, Donghuateng Sports Apparatus Ltd, Beijing, China). All assessments were administered by expert staff from the elite sports schools, and those tests with two repeated trials (which were not requested in the 2-km run, pull-up and plank test) showed test-retest reliability coefficients above *r*_tt_ ≥ 0.80, being fairly adequate for individual measurements.

Before the test session, subjects performed a standardized warm-up consisting of cycling, running and dynamic stretching.

#### Statistical Analysis

All data were analyzed with SPSS (Version 28.0; SPSS Inc., Chicago, IL, United States) and statistical significance was set at *p* ≤ 0.05.

For the evaluation of the anthropometric and motor predictors’ concurrent and discriminative validity, the age influence on the test performances should be considered not only in sports in general(15,33,35,45), but also especially in the track and field throwing disciplines(46,47). Univariate ANOVAs were conducted to check the data set for significant differences of the youth athletes regarding calendar age. As test performances were systematically growing with age in the four adolescent age cohorts (U15 to U18) participating in the study, the calendar age (in months) was partialized out of the results in all predictors by linear regression analyses to avoid confounding effects in the following analysis(27). In the linear regression analyses the test results served as dependent and the age (in months) as independent variables. To allow for comparisons between the different predictors, the residuals of the regressions were standardized by z-values. Henceforth, in all subsequent analyses, only the age-adjusted *z*-values were used.

The concurrent criterion-related validity of the three morphological and twelve physical fitness measures was determined using a bivariate correlation (Pearson) with the throw performance of the discipline-specific standing throw.

The discriminative validity of the three morphological and twelve physical fitness measures was determined using a statistical classification of the athletes by means of a linear discriminant analysis (DA) and a nonlinear neural network MLP. In the first step the four sports served as the dependent grouping variable, whereas the test results were used as an independent variable set. In a second step, the prioritization of the fifteen independent variables was analysed by means of a comparison of one group comprising a single throw discipline with the remaining group of all other three sports together. The aim of this second step was to find out which anthropometric and physical fitness characteristics in particular are the most relevant in each of the four track and field throw disciplines, and to allow for a prioritization of the most important athletes’ characteristics that are specifically valid for each particular throwing discipline. The stepwise DA was based on the “leave-one-out” method. This means the classification of each individual was calculated using a function derived from all other (*n*−1) cases without the single one case that was held out for final classification. Similarly, for the MLP analysis, three subsets were created for (i) training, (ii) validation of the predictive model, and (iii) the final classification of the left-out cases (test sample). Subsequently, the MLP was trained with 70% of all cases, 20% were used for validating the trained network. Finally, the classification was calculated for the test sample of the remaining ten percent of cases. This specific type of leave-out cross validation strategy was repeated ten times so that each case should at least once belong to the hold-out athletes that were finally classified. To quantify the validity of this classification strategy, the percentage of correct hits of the neural network classification was averaged over the ten trials and the mean value was used from there on. Thus, the procedure corresponds to a 10-fold cross-validation. The classification quality of both methods was expressed by the proportion of correct hits that is the percentage of athletes that were assigned as true positives to their own throw discipline. In contrast to the DA, where the different numbers of cases in the two groups are regarded by the a-priori probability and a substantial part of the calculation procedure, in the MLP athletes’ classification the number of cases in the investigated groups should be equivalent. Otherwise, the larger classification groups suck most of the athletes of small groups. To avoid this source of error by group size, in all MLP calculations we assigned the participants of the comparably large group of the remaining three sports randomly into a number of groups that match with the group size of the investigated single sport group. Thus, in shot put and javelin throw we divided the summarized rest group into two, in discus throw into three, and in the hammer throw analysis into 15 groups. In all four analyses the results obtained in the various subgroups were averaged.

## Results

The bivariate correlation analysis revealed a low to medium concurrent validity of the morphological measures and a medium to large concurrent validity of the twelve different physical fitness tests with throwing performance. As Table 2 shows, the criterion-oriented validity of the tests varies between the four throw disciplines.

**Table 1.** Descriptives of the sports-specific throw performances as well as of three morphological and twelve physical fitness characteristics

|  | N | Min | Max | Mean / Standard error |  | Standard deviation |
| --- | --- | --- | --- | --- | --- | --- |
| Shot put standing (m) | 101 | 7,85 | 18,26 | 13,9552 | ,24875 | 2,49988 |
| Hammer throw spinning (m) | 16 | 15,00 | 60,00 | 36,9369 | 4,16362 | 16,65450 |
| Discus throw standing (m) | 63 | 26,00 | 56,00 | 42,8633 | ,69680 | 5,53067 |
| Javelin throw standing (m) | 109 | 15,00 | 50,00 | 37,0424 | ,70217 | 7,33092 |
| Body height (cm) | 256 | 166,00 | 200,00 | 181,5117 | ,40724 | 6,51588 |
| Body weight (kg) | 256 | 52,00 | 145,00 | 87,2500 | 1,11673 | 17,86771 |
| BMI | 289 | 10,00 | 43,20 | 26,2550 | ,29762 | 5,05953 |
| Standing long jump (cm) | 289 | 179,00 | 322,00 | 258,4671 | 1,42778 | 24,27218 |
| Pull-ups (n) | 289 | ,00 | 43,00 | 9,2664 | ,42260 | 7,18420 |
| Plank (s) | 289 | 39,00 | 660,00 | 160,3737 | 4,32420 | 73,51142 |
| Sit and reach (cm) | 289 | 1,00 | 34,00 | 17,6055 | ,34643 | 5,88932 |
| 30-m-sprint (s) | 288 | 3,70 | 6,00 | 4,3650 | ,02306 | ,39129 |
| 60-m-sprint (s) | 281 | 6,60 | 11,39 | 8,0715 | ,04254 | ,71305 |
| Medicine ball throw forward (m) | 289 | 5,00 | 20,55 | 13,3174 | ,13138 | 2,23338 |
| Medicine ball throw backward (m) | 283 | 5,00 | 21,80 | 15,4376 | ,18560 | 3,12221 |
| Hexagonal jump (s) | 289 | 9,70 | 24,30 | 14,1055 | ,12745 | 2,16672 |
| Balance pad (s) | 277 | 10,00 | 120,00 | 50,5343 | 1,28953 | 21,46204 |
| 2-Km-endurance run_(min) | 289 | 7,09 | 13,05 | 9,8523 | ,07696 | 1,30836 |
| Triple jump (m) | 283 | 5,20 | 12,34 | 7,8667 | ,06715 | 1,12964 |

**Table 2.** Criterion-oriented validity coefficients (Pearson correlation, two-sided significance) between test scores and achieved sport-specific throwing performances

|  |  | Shot put standing | Hammer throw spinning | Discus throw standing | Javelin throw standing.<sup>a</sup> |
| --- | --- | --- | --- | --- | --- |
| Body height | Pearson-correlation | $r_{tc} = .10$<br>($p = .347$; $n=92$) | $r_{tc} = .40$<br>($p = .291$; $n=9$) | $r_{tc} = .35$<br>($p = .009$; $n=54$) | $r_{tc} = .20$<br>($p = .047$; $n=101$) |
| Body weight | Pearson-correlation | $r_{tc} = .14$<br>($p = .198$; $n=92$) | $r_{tc} = .15$<br>($p = .695$; $n=9$) | $r_{tc} = .33$<br>($p = .016$; $n=54$) | $r_{tc} = .42$<br>($p < .001$; $n=101$) |
| BMI | Pearson-correlation | $r_{tc} = -.01$<br>($p = .932$; $n=101$) | $r_{tc} = .02$<br>($p = .939$; $n=16$) | $r_{tc} = -.03$<br>($p = .803$; $n=63$) | $r_{tc} = -.04$<br>($p = .657$; $n=109$) |
| Standing long jump | Pearson-correlation | $r_{tc} = .49$<br>($p < .001$; $n=101$) | $r_{tc} = .04$<br>($p = .873$; $n=16$) | $r_{tc} = -.08$<br>($p = .540$; $n=63$) | $r_{tc} = .60$<br>($p < .001$; $n=109$) |
| Pull-ups | Pearson-correlation | $r_{tc} = .64$<br>($p < .001$; $n=101$) | $r_{tc} = .63$<br>($p = .009$; $n=16$) | $r_{tc} = .22$<br>($p = .083$; $n=63$) | $r_{tc} = .31$<br>($p < .001$; $n=109$) |
| Plank | Pearson-correlation | $r_{tc} = .43$<br>($p < .001$; $n=101$) | $r_{tc} = .17$<br>($p = .521$; $n=16$) | $r_{tc} = -.25$<br>($p = .045$; $n=63$) | $r_{tc} = .08$<br>($p = .437$; $n=109$) |
| Sit and reach | Pearson-correlation | $r_{tc} = .58$<br>($p < .001$; $n=101$) | $r_{tc} = -.47$<br>($p = .066$; $n=16$) | $r_{tc} = -.01$<br>($p = .957$; $n=63$) | $r_{tc} = .46$<br>($p < .001$; $n=109$) |
| 30-m sprint | Pearson-correlation | $r_{tc} = -.40$<br>($p < .001$; $n=100$) | $r_{tc} = -.41$<br>($p = .114$; $n=16$) | $r_{tc} = .18$<br>($p = .167$; $n=63$) | $-.413^{**}$ $r_{tc} = -.41$<br>($p < .001$; $n=109$) |
| 60-m sprint | Pearson-correlation | $r_{tc} = -.62$<br>($p < .001$; $n=100$) | $-.268$ $r_{tc} = -.27$<br>($p = .426$; $n=11$) | $r_{tc} = -.20$<br>($p = .119$; $n=62$) | $-.579^{**}$ $r_{tc} = -.58$<br>($p < .001$; $n=108$) |
| Medicine ball throw forward | Pearson-correlation | $r_{tc} = .80$<br>($p < .001$; $n=101$) | $r_{tc} = -.06$<br>($p = .815$; $n=16$) | $r_{tc} = .56$<br>($p < .001$; $n=63$) | $r_{tc} = .53$<br>($p < .001$; $n=109$) |
| Medicine ball throw backward | Pearson-correlation | $r_{tc} = .85$<br>($p < .001$; $n=101$) | $r_{tc} = .78$<br>($p = .005$; $n=11$) | $r_{tc} = .69$<br>($p < .001$; $n=62$) | $r_{tc} = .64$<br>($p < .001$; $n=109$) |
| Hexagonal agility jump | Pearson-correlation | $r_{tc} = -.38$<br>($p < .001$; $n=101$) | $r_{tc} = -.27$<br>($p = .317$; $n=16$) | $r_{tc} = .07$<br>($p = .600$; $n=63$) | $r_{tc} = -.38$<br>($p < .001$; $n=109$) |
| Balance pad stance | Pearson-correlation | $r_{tc} = .46$<br>($p < .001$; $n=92$) | $r_{tc} = -.34$<br>($p = .234$; $n=14$) | $r_{tc} = .06$<br>($p = .621$; $n=62$) | $r_{tc} = .21$<br>($p = .031$; $n=109$) |
| 2-km run | Pearson-correlation | $r_{tc} = -.25$<br>($p = .012$; $n=101$) | $r_{tc} = -.76$<br>($p < .001$; $n=16$) | $r_{tc} = -.03$<br>($p = .812$; $n=63$) | $r_{tc} = .14$<br>($p = .160$; $n=109$) |
| Triple jump | Pearson-correlation | $r_{tc} = .13$<br>($p = .211$; $n=101$) | $r_{tc} = .73$<br>($p = .011$; $n=11$) | $r_{tc} = -.62$<br>($p < .001$; $n=62$) | $r_{tc} = .51$<br>($p < .001$; $n=109$) |

### Classification by linear Discriminant Analysis and nonlinear Neural Network

A total of 8 cases from the four throwing disciplines were excluded from the DA due to missing data. In the first attempt, a DA was calculated with the remaining total sample of *n*=281 cases and a classification rate of 73.0% was obtained. In this analysis, 23 of the 100 shot put athletes (77.0% correct hits), 6 of the included 11 hammer throw athletes (36.4% correct hits), 28 of the 62 discus athletes (45.2% correct hits), and only 19 of the 108 javelin throwers (82.4% correct hits) were assigned erroneously to another discipline. In the second attempt, a cross-validated DA was applied, where each of the 281 athletes was iteratively used as a single hold-out case which has to be solely classified. On the basis of this leave-one-out procedure, 68.7% of all participants were correctly classified and assigned to their own throwing discipline as true positives (Table 3). The best classification result with 82.4% correct hits was again achieved in the javelin throw, where in the cross-validated calculation only 19 out of 108 athletes were assigned as false negatives to another sport (6 to shot put, 2 to hammer throw, and 11 to discus throw). The highest fraction of false negatives was found in the hammer throw (63.7%), mainly due to the erroneous assignment of four (36.4%) youth hammer throwers to the shot put group, and a further three (27.3%) to the javelin group.

**Table 3.** Cross-validated discriminant analysis classification of n = 281 single cases of youth track and field athletes from four throwing disciplines on the basis of 13 performance characteristics (Method: listwise)

| Throwing groups | Predicted throwing discipline |  |  |  |
| --- | --- | --- | --- | --- |
| | Shot put<br>($n$; percent) | Hammer throw<br>($n$; percent) | Discus throw<br>($n$; percent) | Javelin throw<br>($n$; percent) |
| **Shot put**<br>**($n = 100$)** | 72<br>(72.0 %) | 1<br>(1.0 %) | 17<br>(17.0 %) | 10<br>(10.0 %) |
| **Hammer throw**<br>**($n = 11$)** | 4<br>(36.4 %) | 4<br>(36.4 %) | 0<br>(0.0 %) | 3<br>(27.3 %) |
| **Discus throw**<br>**($n = 62$)** | 26<br>(41.9 %) | 1<br>(1.6 %) | 28<br>(45.2 %) | 7<br>(11.3 %) |
| **Javelin throw**<br>**($n = 108$)** | 6<br>(5.6 %) | 2<br>(1.9 %) | 11<br>(10.2 %) | 89<br>(82.4 %) |

Since there were four different sport groups, three linear discriminant functions were established. The first two functions accounted for 92.6% of the total variance and are shown in Figure 1 on the X- and Y-axes . The athletes from the four throw groups are distributed around their respective centroids, which are located on distinct areas of the plot (Functions at group centroids: Shot put, function 1 = 1.10 and function 2 = –0.21; Hammer throw, function 1 = 0.61 and function 2 = –1.37; Discus throw, function 1 = 0.42 and function 2 = 0.74; Javelin throw, function 1 = –1.31 and function 2 = –0.09). The first function (Eigenvalue: 1.15) was the most important, accounting for 78.0% of the variance and was related primarily to the medicine ball throw (backward). The second function (Eigenvalue: 0.22) accounted for 14.6% of the variance and was related especially to the jumping and sprinting performances.

**Figure 1.**
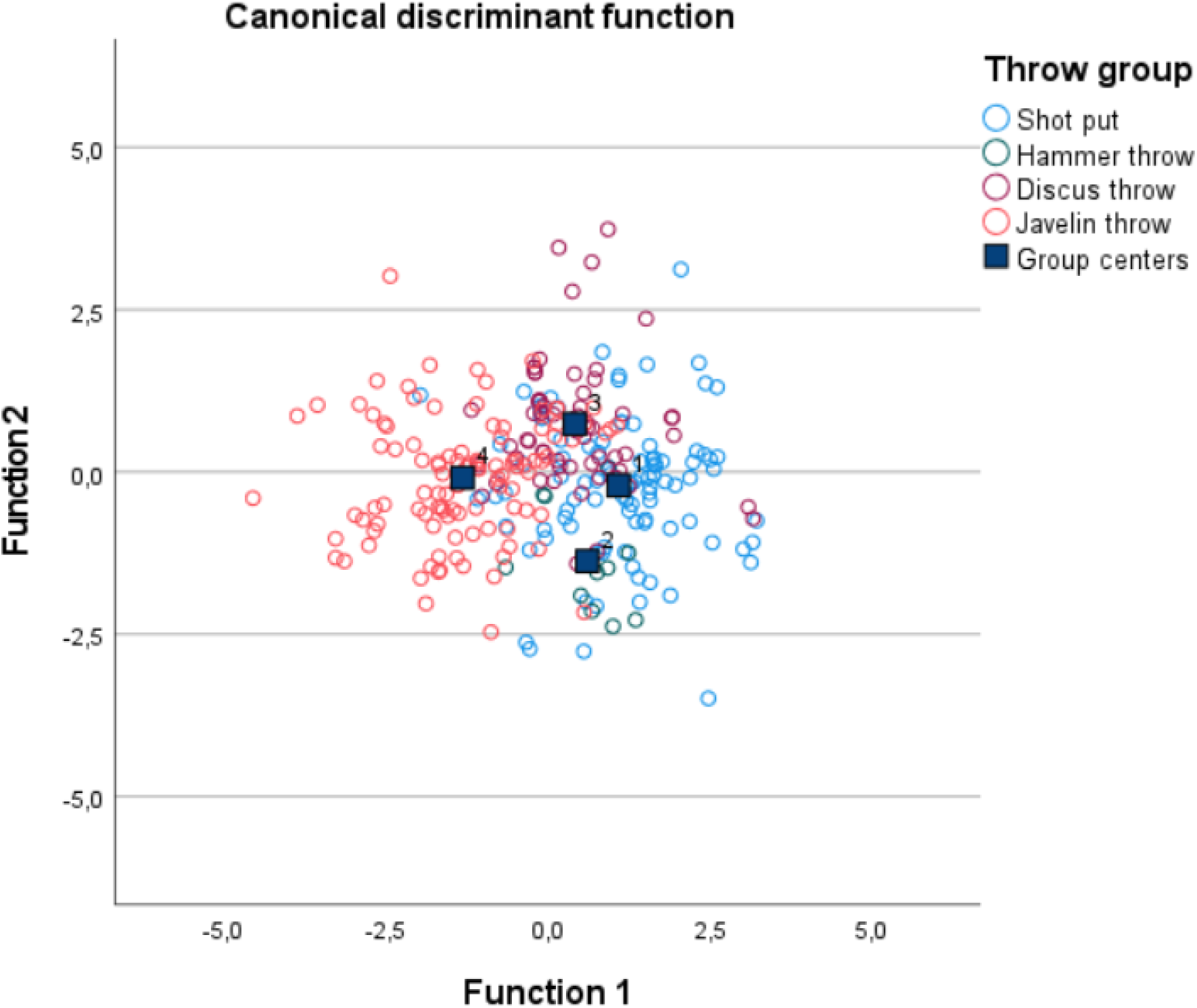
Plot of individual and group differences between the for throw disciplines resulting from one morphological (BMI) and twelve physical fitness tests

### Prioritization of talent characteristics for each throw discipline

In order to analyze the specific talent characteristics that distinguish the athletes of a particular throwing discipline from the athletes of the other three disciplines, a DA was calculated to classify the athletes of that particular discipline from a second group comprising all the other throwing athletes of the three alternative sports. The descriptive statistics of the 15 variables measured in the *N* = 289 male athletes are documented in Table 4. Also shown are the means of age at test, which did not differ systematically between the four groups of throwers (*F*_3;285_ = 1,80; *p* = 0.147).

**Table 4.**
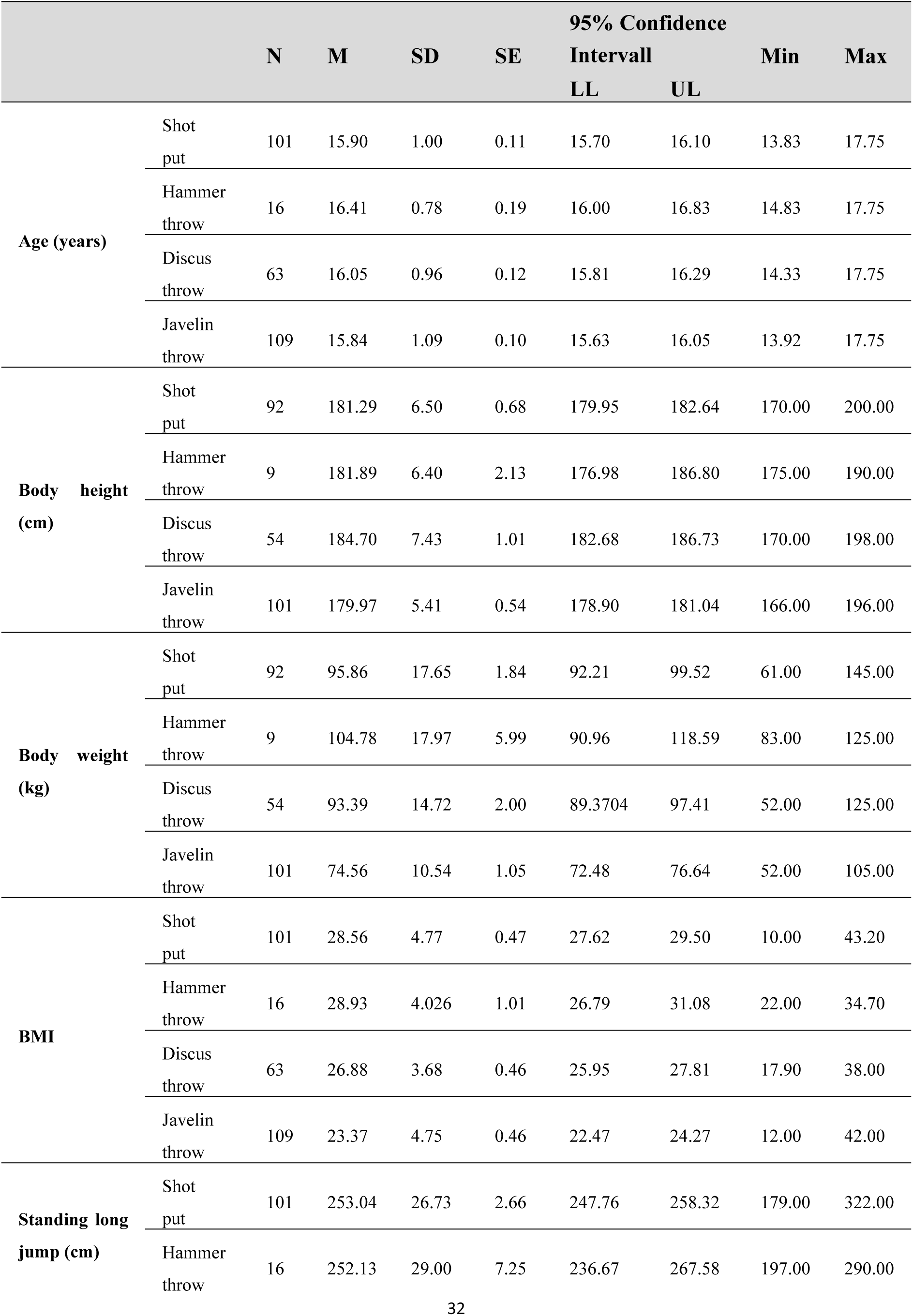
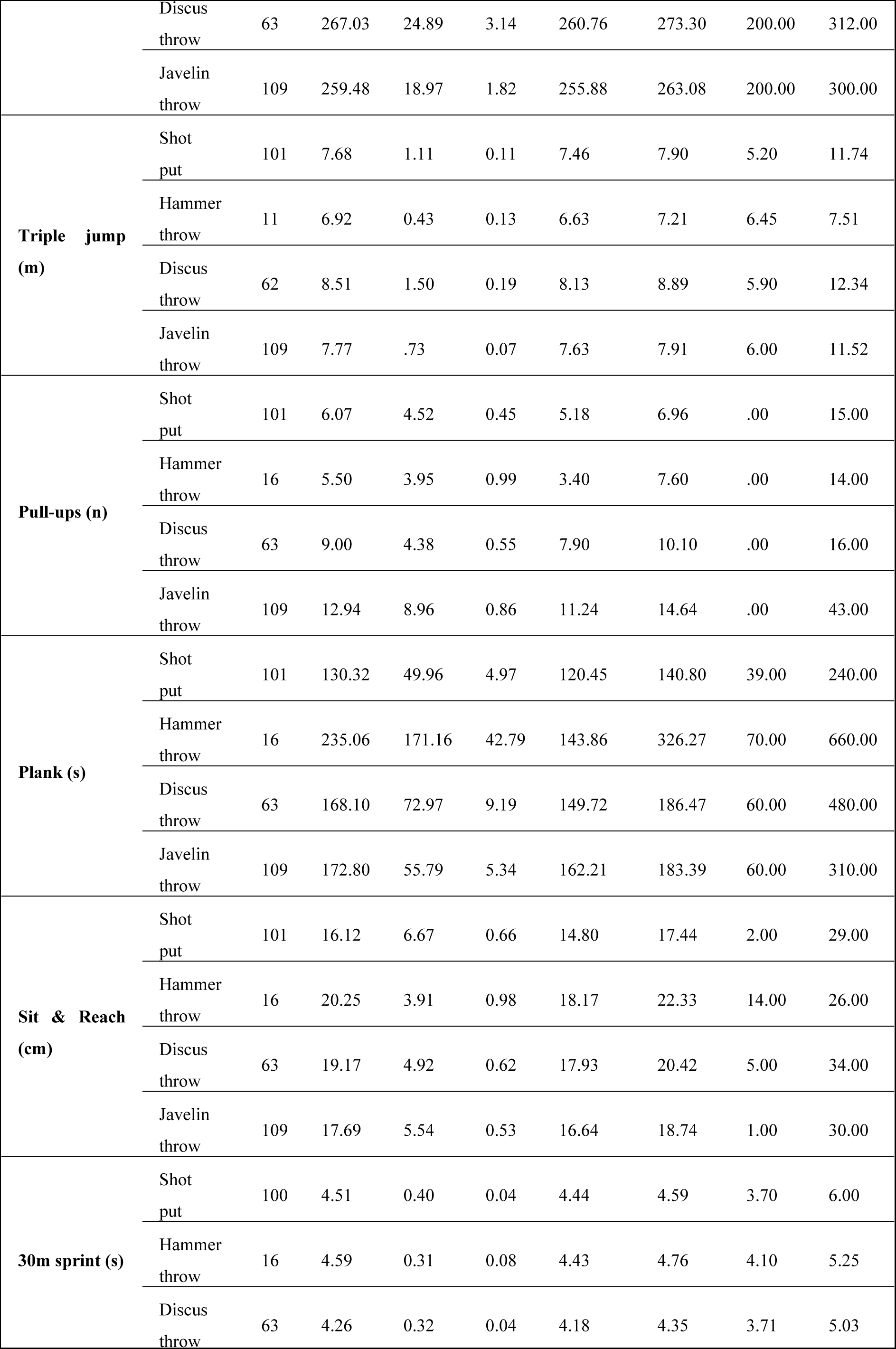
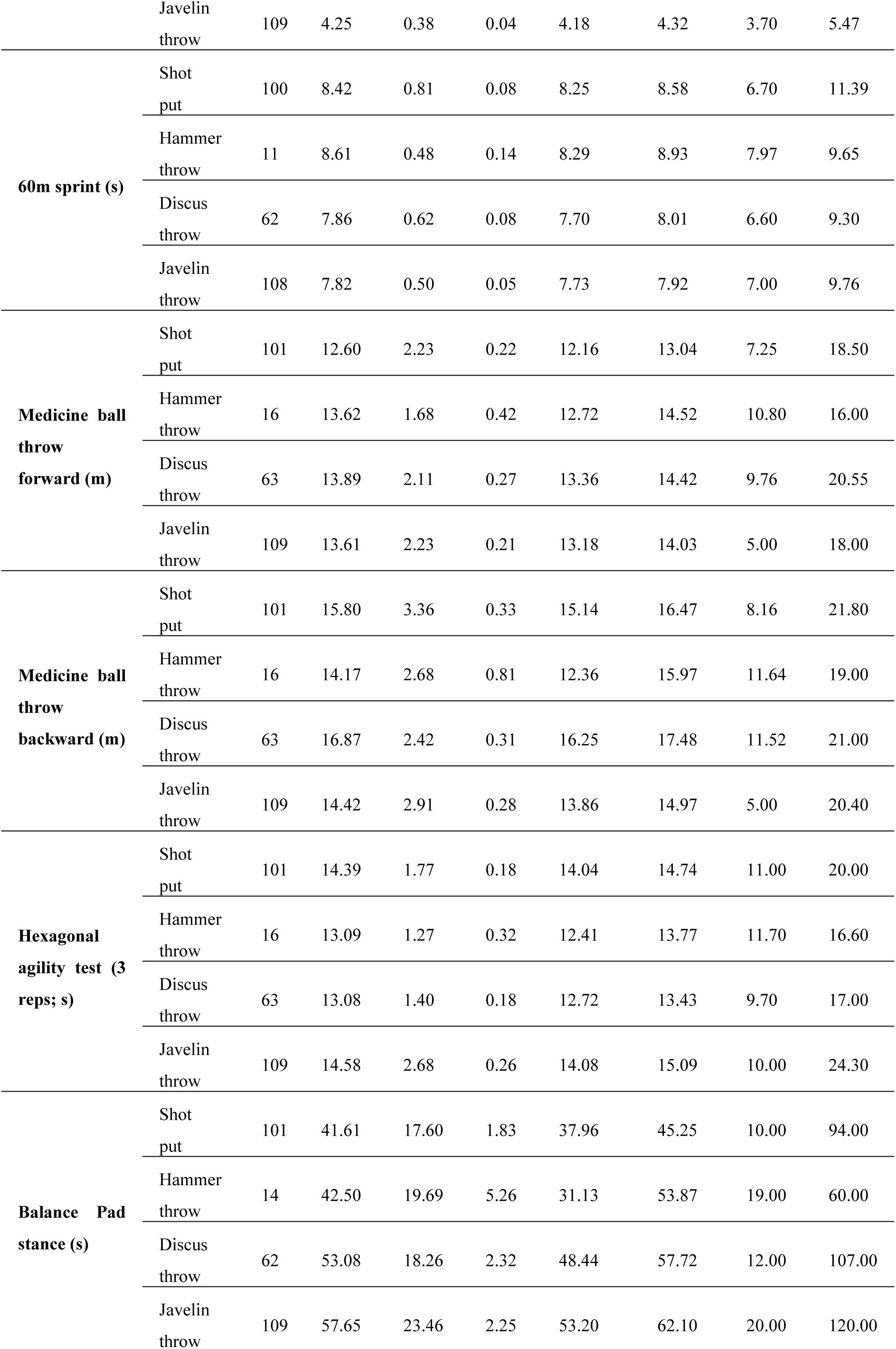
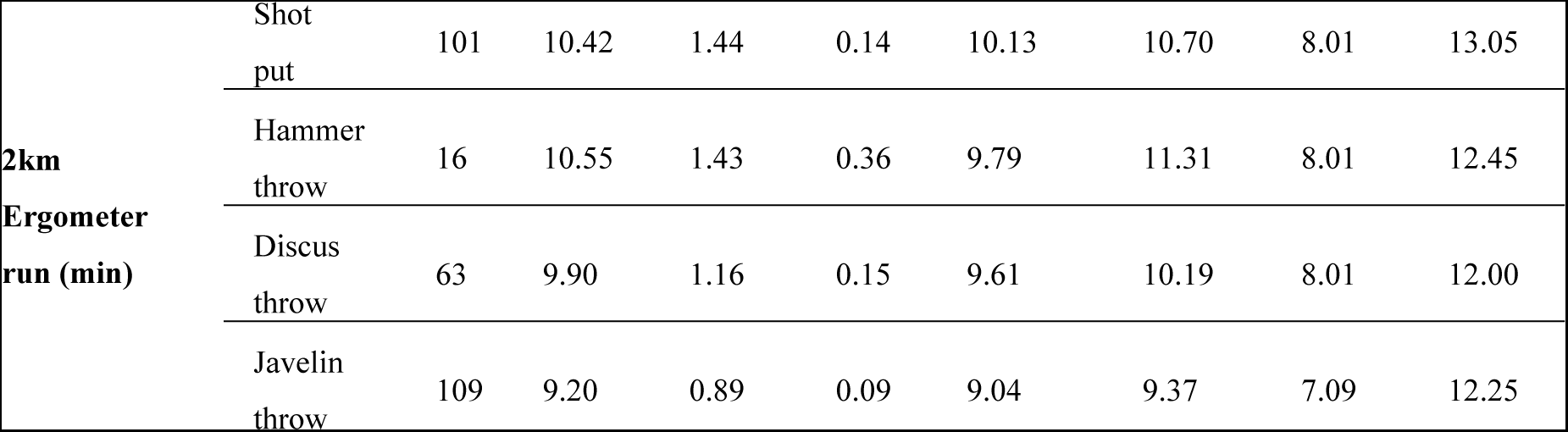
Descriptive data of morphological and physical fitness performance prerequisites of the athletes from four track and field throwing disciplines.

### Shot put

In the stepwise DA of the anthropometric and physical fitness measures, it was shown that the youth shot putters differed most from the rest of all other track and field throwing athletes (see Table 4) in a better medicine ball backward throw performance (discriminant coefficient = 0.70). On the other hand, shot putters performed less well in the medicine ball forward throw (−0.64), 60m sprint (0.38), and the plank test (−0.44). On the basis of these five tests, 69.2% of the 101 shot putters tested could be distinguished from the total group of the 188 athletes from the other throwing disciplines.

In the non-linear MLP analysis (see Fig. 2), the normalized importance of the independent variables in classifying the athletes served as a validity measure to prioritize the anthropometric and physical fitness characteristics which distinguish the participants from the different throwing disciplines. The most relevant motor tests to discriminate shot putters from all other throwing athletes were the forward medicine ball throw (importance: 79.8%, although this test discriminated shot putters from other track and field throwers negatively), and an above average backward throw (importance: 75.8%). In addition, a higher BMI (55.5%), but slower 60m sprint times (63.5%) and also lower standing long jump performance (48.6%) contributed significantly to the correct discrimination of 74.9% of the shot putters from the other throwers.

**Figure 2.**
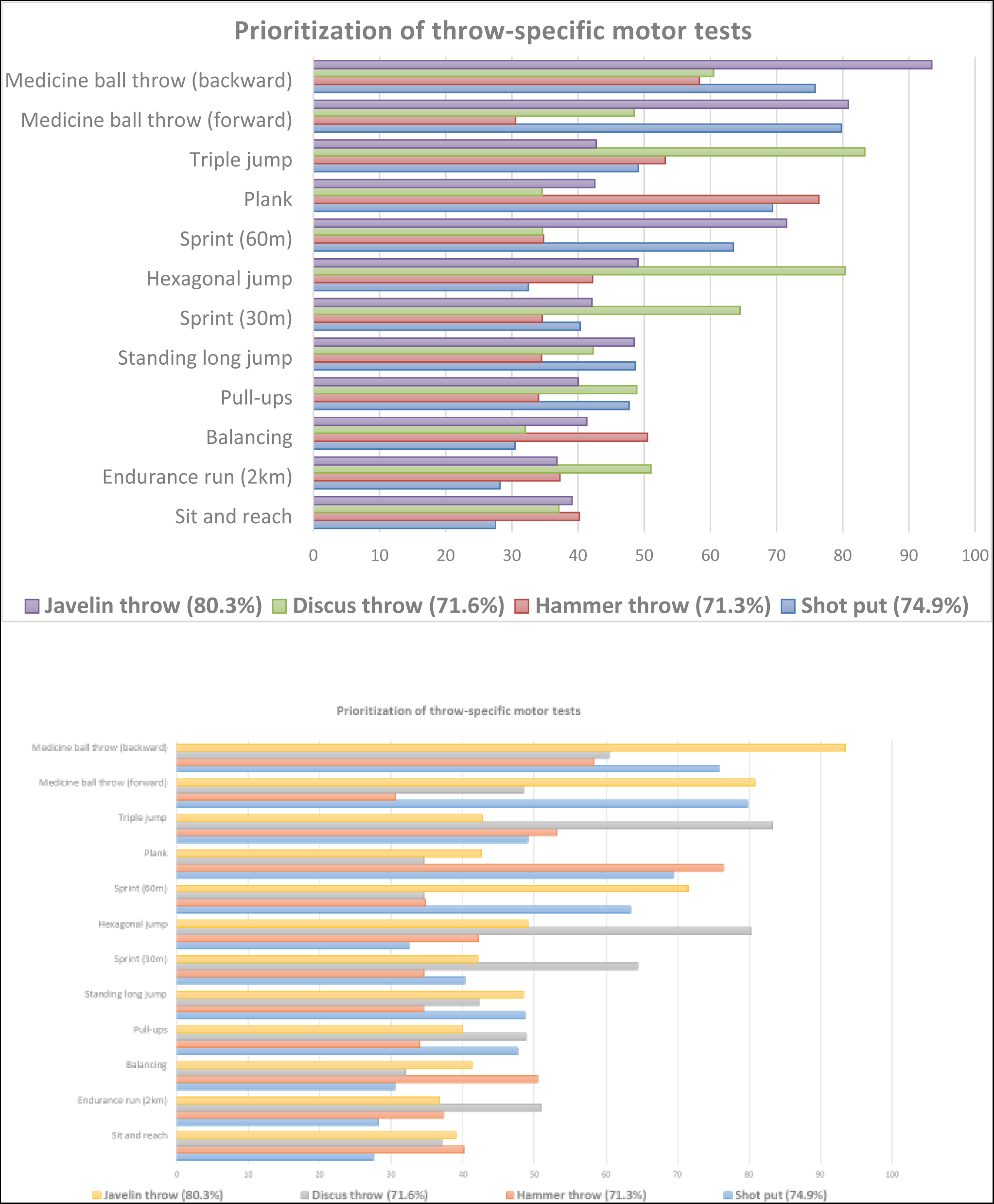
Normalized importance (in percent) of one morphological (BMI) and twelve physical fitness tests in the nonlinear MLP analysis to prioritize each single throw discipline from the remaining total group of all three other sports.

### Hammer throw

The stepwise DA revealed that the hammer throw performers distinguished from the total group of all other youth athletes from the remaining three throwing disciplines mainly by the best core stability performance in the plank test (1.13), a good one-leg balance pad stance duration (0.66) and hexagonal jump agility (−0.26). On the other side, the hammer throwers exhibited a lower triple jump performance (discriminant coefficient = 0.42) and 2-km running endurance (0.34) than their counterparts from the other throwing sports. Nevertheless, 44.4% of the hammer throwers could be successfully classified into their original sport using these five tests. The non-linear MLP analysis corroborated the DA findings, particularly with regard to a systematically lower triple jump performance (normalized importance: 53.2%). On the other hand, the hammer throwers were superior to the other throwing athletes in terms of core stability (plank test importance: 76.4%) and exhibited a higher BMI (importance: 56.7%). Overall, the MLP helped to correctly identify 71.3% of the cases included in the test set and presented to the trained and validated neural network.

### Discus throw

In the discus throwing discipline, the stepwise DA identified three physical fitness tests that distinguished this group from all other youth throwing athletes. So, body height (discriminant coefficient = 0.64), the triple jump distance (discriminant coefficient = 0.70) and the hexagonal jumping agility (−0.37) were superior to all other throwers. On the basis of these three significant tests, the DA could only identify 37.0% of the cases correctly.

The non-linear MLP analysis confirmed the DA results, particularly in the better triple jump (normalized importance: 83.3%), fast 30-m sprint (importance: 64.5%), and better medicine ball backward throw test (importance: 60.5%). On the other hand, the MLP results also confirmed the better hexagonal jumping agility performance (importance: 80.4%), which showed a systematic difference from the other youth throwing athletes. In summary, the MLP was able to distinguish 71.6% of the discus throwers from the remaining other throwing disciplines participants.

### Javelin throw

For the javelin, the stepwise DA produced a comparably long list of nine significant variables, which helped to classify 83.0% of the javelin throwers correctly into their sport. Two anthropometric and seven physical fitness parameters discriminated between the javelin experts and all other participants in track and field throwing disciplines. The highest positive contrast compared to the total group of the other throwers was found in the medicine ball forward throw (discriminant coefficient = −0.64). Furthermore, they showed better sprint performances in the 60-m dash (0.34), the pull-ups test (−0.24), and finally also in the 2-km running endurance trial (0.24). Besides that, a lower BMI (0.35) was diagnosed, mainly due to a lower body weight (0.39). In contrast to these six performance characteristics, javelin throwers were less proficient than their counterparts from the group of the other three throwing disciplines especially in the backward medicine ball throw (0.65), but also in the standing long jump (0.43) and the jumping agility, measured by the hexagonal jump test (−0.23).

The non-linear MLP analysis confirmed the DA results to a large extent, as the performance gap to the total group of other athletes was systematic in the poorer medicine ball backward throw (normalized importance: 93.5%). In addition, lower body weight (85.3%) and a lower BMI (56.5%), better 60-m sprint performance (71.5%), and an above average jumping agility in the hexagonal test (49.1%) also contributed to the correct identification of 80.3% of the javelin throw specialists.

## Discussion

The main objective of this research was to discriminate between youth track and field athletes from more than 30 Chinese elite sports schools - many of whom will contribute to the next generation of elite senior athletes—belonging to four throwing disciplines sports. Therefore, a generic test battery of 15 anthropometric and motor performance characteristics was administered to 289 adolescent elite athletes. It should be highlighted that in this study both linear and non-linear statistical methods were used in parallel to identify the most relevant throwing performance characteristics for each of the four throwing disciplines and to confirm the results of each method in turn. The main findings included the high discriminative validity of the generic test battery that allowed the cross-validated correct assignment of 68.2% of cases using linear discriminant analysis, and an impressive rate of 72.2% using the non-linear neural network tool MLP. The high quality of the roughly three-quarters correct discriminations between the four groups of adolescent shot putters, hammer, discus and javelin throwers can be described as good, and is underlined by the fact that in this study a very homogeneous sample of throwing athletes from track and field sport only was investigated. The worth of this finding is also evident when compared to the cross-validated results of 71.3% correct assignments reported by Zhao et al. (25)in a comparable study in a Chinese elite sport school in Shanghai on athletes from six much more heterogeneous sports (basketball, fencing, judo, swimming, table tennis, volleyball). Better results can be obtained if one refrains from checking the classification results by means of a cross-validation procedure, as can be seen in the study by Pion et al. (32) for a selection of nine very different sports (badminton, basketball, gymnastics, handball, judo, soccer, table tennis, triathlon, and volleyball). The 100% classification reached by Pion et al.(41) in a discriminant analysis in the three more homogeneous martial arts disciplines judo, karate, and taekwondo also underlines the importance of a cross-check procedure using a leave-one-out strategy or even a larger hold-out group. Our results are also promising when compared to the 88.0% classification rate in a DA of Leone et al. (39) in the heterogeneous total group of figure skaters, swimmers, tennis, and volleyball players, as well as the 85.2% correct hits reported by Opstoel et al. (40) for a DA in ball sports, dance, gymnastics, martial arts, racket sports, and swimming athletes. In summary, our study result of almost three-quarters correct classifications by means of a nonlinear neural network in throwing athletes cannot be directly compared with the results of research groups that calculated the predictive accuracy only for the original sample, including all members of the total groups. Nevertheless, despite the introduction of hold-out cases in the classification for cross-check purposes our results are still very satisfactory. This holds true in all four the disciplines and especially in javelin throw, whereas the linear DA analysis failed particularly in the identification of hammer and discus throw athletes.

The best classification results with 82.4% correct hits in the DA were obtained in the javelin throw. The high predictive accuracy is mainly based on the standing long jump and triple jump tests as well as the two sprint and the two medicine ball throw performances, which is in line with Bouhlel, et.al.(7) and Schleichardt, et.al.(10), who attributed this finding to the relevance of explosive strength abilities in this particular sport, and where body dimensions (e.g. body weight) do not play such an important role as in the discus throw and shot put disciplines (2–4). Similarly, in the neural network analysis using the MLP tool and the use of the ten-percent-hold-out strategy, we discovered that 80.3% of the javelin throwers were assigned correctly to their original sport. Although, that we expected that the non-linear neural network method to give a higher classification rate than the linear discriminant analysis, the equality with the DA result at a comparably high level is convincing.

In the shot put and the discus throw groups the DA produced correct classification results of 72.0%, and 45.2% respectively. As mentioned above, especially the anthropometric characteristic of body height plays an important role in shot put and discus throw athletes, as a large height allows for a high take-off height and, in combination with the longer arms cause a long acceleration trajectory of the heavy competition device(48). In addition to the similar anthropometric performance prerequisites, the performances of the throwers from these two disciplines in the medicine ball throws (forward and backward) and the above-average jumping abilities also have an important influence on the over-all performance in the sports-specific throws.

In the group of hammer throw athletes, there were still significant deficits in the prediction accuracy of the linear DA, where about two thirds of the hammer throw participants being not correctly identified. Although that the nonlinear MLP method produced far better predictions, this finding may be attributed to the much smaller size of this sport-specific group. Such a small fraction of participants makes it difficult to identify a typical sport-specific performance prerequisites profile, which can be distinctively separated from the comparatively large number of shot put, discus and javelin cases. Thus, the matching of the group sizes used in the MLP procedure seems to be more adequate than the use of the a-priori probability in the DA analysis. In the past, talent identification for certain sports was based primarily on sport-specific tests, which made it difficult not only to compare results between different sports, but also to draw conclusions about the possibility of transferring an athlete from a donor-sport to a recipient sport, as has become increasingly important in elite sport over the last two decades(49). Also, such sport-specific tests pose the problem that athletes from one sport who are not familiar with the specific techniques and skills of an alternate sport cannot perform tests other than those specific to their own sport with a reliable and valid personal result. It is therefore crucial that both talent development and talent transfer programs consist of a multidisciplinary mix of anthropometric, motor, psychological, or physiological tests of low to medium specificity. Such complex test profiles allow a more complete assessment of each athlete, as well as between-sports comparisons, and cross-disciplines transfer campaigns(32,41,50). In this sense, the results of this study provide substantial information about the validity of the anthropometric and motor performance tests for discriminating young athletes from different inter-related sports disciplines. Furthermore, we are provided with evidence for the applicability of the different measures for talent transfer in the four track and field throw disciplines investigated. The combination of linear DA and nonlinear MLP neural networks is a fruitful approach for resolving the problem of talent education as well as talent transfer, especially when it is assumed that different types of talent patterns exist in the make-up of promising youngsters, which may lead to the same performance outcome in later developmental stages of an elite sports career(50–53). As both methods on the basis of the age-corrected data set led to an at least satisfactory overall rate of 68.2% (DA) and an even more convincing rate of 72.2% (MLP) correct assignments to the four track and field throwing disciplines, both classification methods seem to establish a broadly comparable relationship between the identified patterns of the 15 predictors and the four sport categories. The most relevant differences between the results of the two analytical models was found in regard to the hammer and the discus throw, as the DA correctly identified only 45.2% of the youth discus and 36.4% of the hammer throwers.

Our study has several limitations. The first limitation is the relatively small sample size, especially in hammer throw (*n*=16 athletes, but only eleven cases with complete data sets), which may be a consequence of the high level of specialization of Chinese elite sport school athletes in this technically demanding throwing discipline. All applicants are subject to strict sport-specific selection and training at an early age, which implies that the highly specialized hammer throw athletes were less numerous in the age group of 14–18-year-old male athletes studied. The fact that hammer throwers cannot participate in decathlon events may also limit participation in this particular discipline. A second limitation is the focus solely on male youth athletes only. In addition to the even smaller number of female youth athletes in the Chinese elite sports schools, the decision to select the male athletes was also a result of the complexity of the study due to the different influence of the gender-specific athletic make-up on the sport-specific performance of male and female youth athletes in the four throwing disciplines studied.

## Conclusion

The results of this study show that in 14-18 years old male and sub-elite cadre athletes from more than 30 Chinese elite sport schools, the differences between sports on a battery of generic anthropometric and motor performance tests allow one to distinguish more than two out of three young athletes’ talents according to their individual sports provenience, regardless of the classification method used. Furthermore, the overall accuracy of the talent classification in the Chinese elite youth athletes is at the level found in European studies. In order to make such comparisons between sports, it is necessary that talent identification programs consist of a multidisciplinary mix of anthropometric and physical fitness tests of low specificity, which in the future could perhaps be enriched with additional psychological or physiological tests. The parallel use of linear and non-linear classification methods to identify the most relevant talent characteristics of each of the four throwing disciplines and the use of hold-out cases for the purpose of cross-validation also enhanced the quality of the results. With regard to the relevance of the different sport-specific athletes’ characteristics for talent transfer campaigns in the practical area of track and field sports, the applied talent classification strategies underlined the importance of superior stature measures and jumping power primarily in the discus throw. Shot putters and hammer throwers were found to have better arm strength abilities, whereas javelin throwers were found to present superior running sprint performances. All participants, except the small group of hammer throwers, showed high levels of explosive arm power in the medicine ball throwing exercises. This was particularly obvious for the shot put and discus throwers. The limitations of the relatively small sample size in hammer throw and the focus on male athletes only require further investigation in the talent composition of elite youth sport cadres. In addition, a greater variety of motor skills tests, including tests of coordination and technical ability, would have been useful in the assessment of young athletes.

## Author Contributions

KZ was responsible for data collection, financial support and test organization as well as revising the manuscript. YZ was responsible for data collection. AH was responsible for data analysis. MS was responsible for writing draft of the manuscript.

## References

1. Crandall CG. Alterations in human baroreceptor reflex regulation of blood pressure following 15 days of simulated microgravity exposure. Citeseer; 1993.

2. Zaras N, Stasinaki A-N, Terzis G. Biological determinants of track and field throwing performance. J Funct Morphol Kinesiol. 2021;6(2):40.

3. Carter JEL, Aubry SP, Sleet DA. 5. Somatotypes of Montreal Olympic Athletes. In: Physical structure of Olympic athletes. Karger Publishers; 1982. p. 53–80.

4. Morrow Jr JR, Disch JG, Ward PE, Donovan 3rd TJ, Katch FI, Katch VL, et al. Anthropometric, strength, and performance characteristics of American world class throwers. J Sports Med Phys Fitness. 1982;22(1):73–9.

5. Łysoń-Uklańska B, Błażkiewicz M, Kwacz M, Wit A. Muscle force patterns in lower extremity muscles for elite discus throwers, javelin throwers and shot-putters–a case study. J Hum Kinet. 2021;78(1):5–14.

6. Terzis G, Georgiadis G, Vassiliadou E, Manta P. Relationship between shot put performance and triceps brachii fiber type composition and power production. Eur J Appl Physiol. 2003;90:10–5.

7. Bouhlel E, Chelly MS, Tabka Z, Shephard R. Relationships between maximal anaerobic power of the arms and legs and javelin performance. J Sports Med Phys Fitness. 2007;47(2):141.

8. Stone MH, Sanborn K, O’Bryant HS, Hartman M, Stone ME, Proulx C, et al. Maximum Strength-Power-Performance Relationships in Collegiate Throwers. J Strength Cond Res. 2003;17(4):739–45.

9. Zaras N, Spengos K, Methenitis S, Papadopoulos C, Karampatsos G, Georgiadis G, et al. Effects of strength vs. ballistic-power training on throwing performance. J Sports Sci Med. 2013;12(1):130.

10. Schleichardt A, Badura M, Lehmann F, Ueberschär O. Comparison of force-velocity profiles of the leg-extensors for elite athletes in the throwing events relating to gender, age and event. Sport Biomech. 2021;20(6):720–36.

11. Caughey R, Thomas C. Variables Associated with High School Shot Put Performance. Int J Exerc Sci. 2022;15(6):1357–65.

12. Jha P, Nuhmani S, Kapoor G, Al Muslem WH, Joseph R, Kachanathu SJ, et al. Efficacy of core stability training on upper extremity performance in collegiate athletes. J Musculoskelet Neuronal Interact. 2022;22(4):498.

13. Okada T, Huxel KC, Nesser TW. Relationship between core stability, functional movement, and performance. J Strength Cond Res. 2011;25(1):252–61.

14. Kim H, Lee Y, Shin I, Kim K, Moon J. Effects of 8 weeks’ specific physical training on the rotator cuff muscle strength and technique of javelin throwers. J Phys Ther Sci. 2014;26(10):1553–6.

15. Collins R, Collins D, MacNamara Á, Jones MI. Change of plans: an evaluation of the effectiveness and underlying mechanisms of successful talent transfer. J Sports Sci. 2014;32(17):1621–30.

16. MacNamara Á, Collins D. Second chances: investigating athletes’ experiences of talent transfer. PLoS One. 2015;10(11):e0143592.

17. Wormhoudt R, Savelsbergh GJP, Teunissen JW, Davids K. The athletic skills model: optimizing talent development through movement education. Routledge; 2017.

18. Bazyler CD, Mizuguchi S, Harrison AP, Sato K, Kavanaugh AA, DeWeese BH, et al. Changes in muscle architecture, explosive ability, and track and field throwing performance throughout a competitive season and after a taper. J Strength Cond Res. 2017;31(10):2785–93.

19. Sakamoto A, Kuroda A, Sinclair PJ, Naito H, Sakuma K. The effectiveness of bench press training with or without throws on strength and shot put distance of competitive university athletes. Eur J Appl Physiol. 2018;118:1821–30.

20. Young KP, Haff GG, Newton RU, Gabbett TJ, Sheppard JM. Assessment and monitoring of ballistic and maximal upper-body strength qualities in athletes. Int J Sports Physiol Perform. 2015;10(2):232–7.

21. Horst F, Janssen D, Beckmann H, Schöllhorn WI. Can individual movement characteristics across different throwing disciplines be identified in high-performance decathletes? Front Psychol. 2020;11:2262.

22. Matthie JR, Withers PO, Van Loan MD, Mayclin PL. Development of a commercial complex bio-impedance spectroscopic (CBIS) system for determining intracellular water (ICW) and extracellular water (ECW) volumes. In: Proceedings of 8th International Conference on Electrical Bio-impedance. 1992. p. 203–5.

23. Kinugasa T. Targeting Tokyo 2020 and beyond: The Japanese TID model. In: Proceedings of the Conference on Talent Identification–Identifying Champions, Doha: Aspire Academy. 2014.

24. Pion J. The Flemish Sports Compass. From Sports Orientation to Elite Performance Prediction. Ghent University; 2015.

25. Zhao K, Hohmann A, Chang Y, Zhang B, Pion J, Gao B. Physiological, anthropometric, and motor characteristics of elite Chinese youth athletes from six different sports. Front Physiol. 2019;10(APR).

26. Müller L, Müller E, Kornexl E, Raschner C. The relationship between physical motor skills, gender and relative age effects in young Austrian alpine ski racers. Int J Sports Sci Coach. 2015;10(1):69–85.

27. Hohmann A, Siener M. Talent identification in youth soccer: Prognosis of U17 soccer performance on the basis of general athleticism and talent promotion interventions in second-grade children. Front Sport Act Living. 2021;3:625645.

28. Fernandez-Fernandez J, Ulbricht A, Ferrauti A. Fitness testing of tennis players: How valuable is it? Br J Sports Med. 2014;48(Suppl 1):i22–31.

29. Siener M, Faber I, Hohmann A. Prognostic validity of statistical prediction methods used for talent identification in youth tennis players based on motor abilities. Appl Sci. 2021;11(15):7051.

30. Faber IR, Bustin PMJ, Oosterveld FGJ, Elferink-Gemser MT, Nijhuis-Van der Sanden MWG. Assessing personal talent determinants in young racquet sport players: a systematic review. J Sports Sci. 2016;34(5):395–410.

31. Siener M, Hohmann A. Talent orientation: The impact of motor abilities on future success in table tennis. Ger J Exerc Sport Res. 2019;49:232–43.

32. Pion J, Segers V, Fransen J, Debuyck G, Deprez D, Haerens L, et al. Generic anthropometric and performance characteristics among elite adolescent boys in nine different sports. Eur J Sport Sci. 2015;15(5):357–66.

33. Meylan C, Cronin J, Oliver J, Hughes M. Talent identification in soccer: The role of maturity status on physical, physiological and technical characteristics. Int J Sports Sci Coach. 2010;5(4):571–92.

34. Unnithan V, White J, Georgiou A, Iga J, Drust B. Talent identification in youth soccer. J Sports Sci. 2012;30(15):1719–26.

35. Höner O, Votteler A. Prognostic relevance of motor talent predictors in early adolescence: A group-and individual-based evaluation considering different levels of achievement in youth football. J Sports Sci. 2016;34(24):2269–78.

36. Buekers M, Borry P, Rowe P. Talent in sports. Some reflections about the search for future champions. Mov Sport Sci Mot. 2015;(88):3–12.

37. Williams AM, Franks AM. Talent identification in soccer. Sport Excercise Inj. 1998;4(4):159–65.

38. Williams AM, Reilly T. Talent identification and development in soccer. J Sports Sci. 2000;18(9):657–67.

39. Leone M, Lariviere G, Comtois AS. Discriminant analysis of anthropometric and biomotor variables among elite adolescent female athletes in four sports. J Sports Sci. 2002;20(6):443–9.

40. Opstoel K, Pion J, Elferink-Gemser M, Hartman E, Willemse B, Philippaerts R, et al. Anthropometric characteristics, physical fitness and motor coordination of 9 to 11 year old children participating in a wide range of sports. PLoS One. 2015;10(5):e0126282.

41. Pion J, Fransen J, Lenoir M, Segers V. The value of non-sport-specific characteristics for talent orientation in young male judo, karate and taekwondo athletes. Arch Budo. 2014;10(1):147–54.

42. Bullock N, Gulbin JP, Martin DT, Ross A, Holland T, Marino F. Talent identification and deliberate programming in skeleton: Ice novice to Winter Olympian in 14 months. J Sports Sci. 2009;27(4):397–404.

43. Stewart AD, Marfell-Jones M, Olds T, Ridder HD. International Standards for Anthropometric Assessment. Wellington, New Zealand: International Society for the Advancement of Kinanthropometry. 2011.

44. Hawes MR, Martin AD. Human Body Composition In: Kinanthropometry and exercise physiology laboratory manual: Tests, procedures and data second edition. Volume 1: Anthropometry. London, Routledge; 2001.

45. Höner O, Leyhr D, Kelava A. The influence of speed abilities and technical skills in early adolescence on adult success in soccer: A long-term prospective analysis using ANOVA and SEM approaches. PLoS One. 2017;12(8):e0182211.

46. Figueiredo LS, Silva DG da, Oliveira BHG, Ferreira AG, Gantois P, Fonseca F de S. Relative age effects in elite Brazilian track and field athletes are modulated by sex, age category, and event type. Mot Rev Educ Física. 2021;27.

47. Redondo JC, Fernández-Martínez E, Izquierdo JM. Effect of relative age on the throwing disciplines of Spanish participants in the national athletics technification plan. Cuad Psicol del Deport. 2019;19(3):156–67.

48. Mastalerz A, Sadowski J. Variability of Performance and Kinematics of Different Shot Put Techniques in Elite and Sub-Elite Athletes–A Preliminary Study. Int J Environ Res Public Health. 2022;19(3):1751.

49. Teunissen JWAJW, Ter Welle SS, Platvoet SSWJ, Faber I, Pion J, Lenoir M. Similarities and differences between sports subserving systematic talent transfer and development: The case of paddle sports. J Sci Med Sport. 2021;24(2):200–5.

50. Pion J, Hohmann A, Liu T, Lenoir M, Segers V. Predictive models reduce talent development costs in female gymnastics. J Sports Sci. 2017;35(8):806–11.

51. Philippaerts R, Coutts A, Vaeyens R. Physiological perspectives of the identification and development of talented performers in sport. In: Talent identification and development-the search for sporting excellence. H&P Druck; 2008. p. 49–68.

52. Pfeiffer M, Hohmann A. Applications of neural networks in training science. Hum Mov Sci. 2012;31(2):344–59.

53. Till K, Jones BL, Cobley S, Morley D, O’Hara J, Chapman C, et al. Identifying talent in youth sport: a novel methodology using higher-dimensional analysis. PLoS One. 2016;11(5):e0155047.

